# Sex change every year: A unique reproductive strategy of the stony coral, *Fimbriaphyllia* (*Euphyllia*) *ancora*

**DOI:** 10.1101/2023.12.27.573442

**Authors:** Shinya Shikina, Pin-Hsuan Tsai, Yi-Ling Chiu, Ching-Fong Chang

**Affiliations:** Institute of Marine Environment and Ecology, National Taiwan Ocean University, Keelung, Taiwan; Center of Excellence for the Oceans, National Taiwan Ocean University, Keelung, Taiwan; Department of Aquaculture, National Taiwan Ocean University, Keelung, Taiwan

**Keywords:** Coral, Sex change, *Fimbriaphyllia ancora*, Annual sex change

## Abstract

The present study documents a unique reproductive strategy of the colonial stony coral, *Fimbriaphyllia ancora,* during observations spanning 8 years. Of 26 colonies monitored at Nanwan Bay, southern Taiwan, about 70% changed their sexes every year, i.e., colonies that were males two years ago became females last year, and changed back to males this year. Apparently, the remaining 30% were permanently male or female. Sex-change and non-sex-change colonies were growing in close proximity or even side-by-side, suggesting that this sex change phenomenon is not driven by environmental factors. No significant differences were found in colony size between sex-change and non-sex-change colonies, suggesting that the sex change strategy may be related to intrinsic factors, e.g., age or genetics. Histological analysis showed that female-to-male sex change occurs 4-5 months after spawning, whereas male-to-female sex change occurs 0-3 months after sperm release. We propose that this unique strategy may increase success of sexual reproduction of sessile, colonial corals.

## Introduction

Sex change is a phenomenon in which an individual reproduces as one sex during a given reproductive season, and as the other sex subsequently (1). Sex change is classified as protandrous (from male to female), protogynous (from female to male), or serial/bidirectional (2-5). Sex change has been observed in various animals and plants, and is considered a reproductive strategy to increase odds of successful propagation (4). Factors triggering sex change, such as size, age, and social factors (the disappearance of a male or female from a group) differ among species (5,6).

Stony corals are the keystone species of coral reef ecosystems, creating complex 3D structures that support the highest marine biodiversity on the planet (7,8). Sexual reproduction of more than 400 coral species (about one-third of existing species) has already been documented over the past 4 decades, and it has been shown that ∼70% of corals are hermaphroditic and ∼30% are gonochoric (9). Sex change has been observed in some species belonging to the Family Fungiidae, which is composed of free-living, non-sessile species (10,11). These species are males when they are small and become females after reaching a certain size, i.e., protandrous sex change (10,11). Bidirectional sex change has also been observed in some fungiid corals (10-12).

*Fimbriaphyllia ancora* (Family Euphyllidae) is a reef-building coral widely distributed in Indo-Pacific tropical and subtropical waters (13,14). This is a colonial species forming a dome or cushion-shaped colony, and is typified by its flabello-meandroid skeleton and tentacles with anchor-like (or hammer-like) tips **(Fig. 1 A and B)**. Multiple colour variants, including green, purple, and orange, are present, making them one of the most popular species in the aquarium ornamental industry (15). This animal is described as a gonochoric species and an annual spawner (16,17). Distinct development of ovaries and testes can be observed in female and male colonies, respectively, nearing the reproductive season **(Fig. 1 C and D)**.

**Figure 1.**
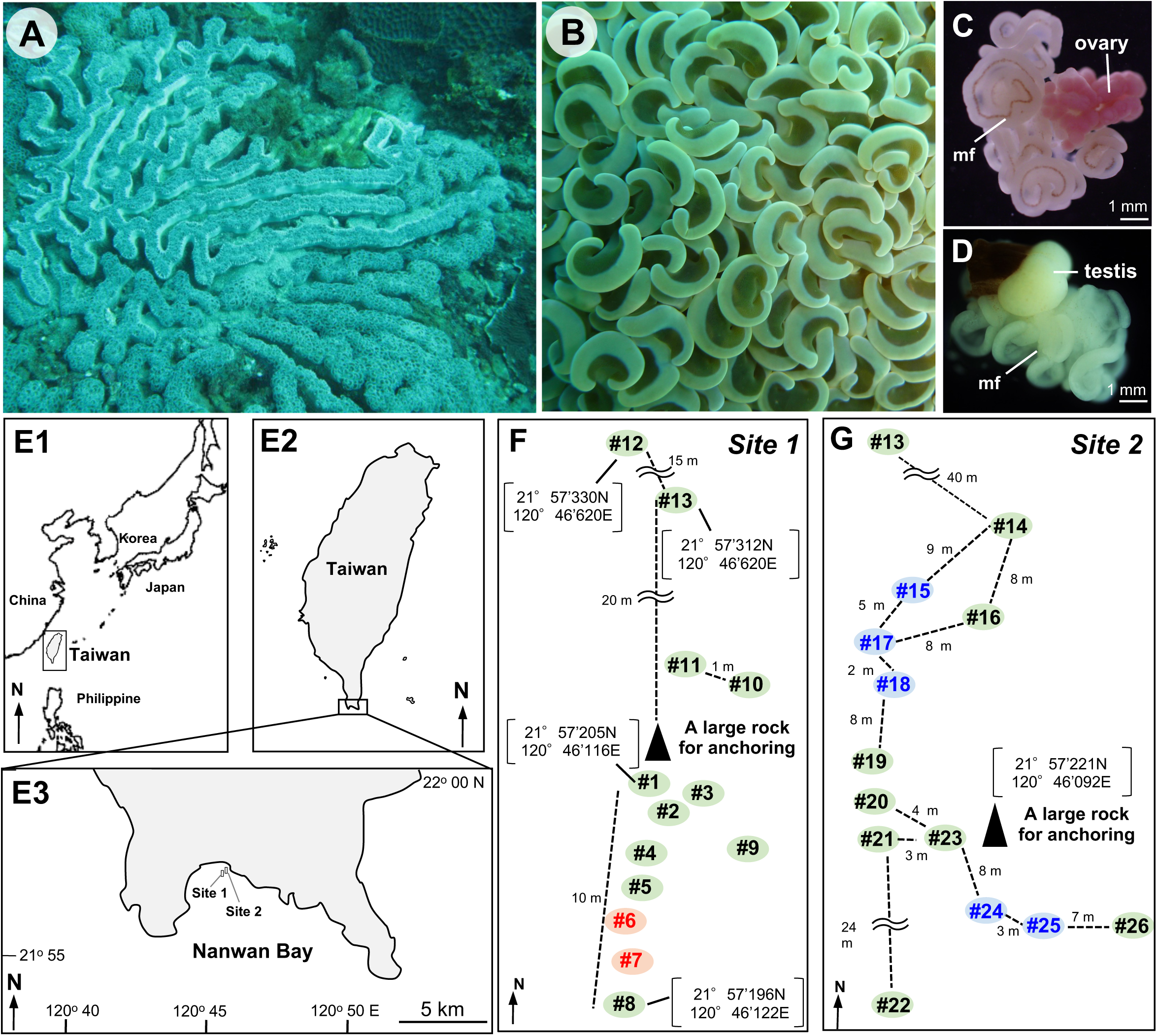
External appearance of *Fimbriaphyllia ancora* and colony locations in Nanwan Bay, Taiwan. **A**. External appearance of an *F. ancora* colony from Nanwan Bay, Taiwan. **B**. External appearance of tentacles of the *F. ancora* colony. The flabello-meandroid skeleton and anchor-like tentacles typify *F. ancora*. **C**. Representative micrograph of an ovary with a mesenterial filament (mf). **D**. Representative micrograph of a testis with a mesenterial filament. **E1-E3**. Maps of survey/sampling locations in Nanwan Bay, southern Taiwan. The map **E3** shows the approximate location of Sites 1 and 2, where *F. ancora* colonies were collected. **F** and **G**. Illustrations showing the approximate locations of *F. ancora* colonies in Site 1 (**F**) and Site 2 (**G**). Numbers (#1-26) indicate colony IDs. Approximate distances of sex-change colonies (letters in black), non-sex-change female colonies (letters in red), and non-sex-change male colonies (letters in blue) are shown.

In 2010 at Nanwan Bay, southern Taiwan, our ecological survey found two sites with many *F. ancora* colonies **(Fig. 1 E1-E3, F, and G)**. We labeled 11 *F. ancora* colonies at one site (Site 1) to understand their reproductive characteristics, and sampled them several times, commencing a few months before spawning, and analyzed sex and gametogenesis histologically. In February 2011, about a year later, we resampled and analyzed the labeled colonies. To our surprise, we found that 9 of 11 labeled colonies had switched sex. At that time, sex change in corals had only been reported in some fungiid corals (10,11). Sex change in other families of corals, especially sessile colonial corals, had not been reported. Then, we hypothesized that *F. ancora* changes its sex at the colony level, and sought to verify this hypothesis by long-term surveys spanning multiple reproductive seasons.

Here we present a unique reproductive strategy, annual sex change of *F. ancora*. Additionally, for the first time in corals, we present histological evidence of the timing and processes of that sex change.

## Results

### Discovery of annual sex change in F. ancora

After noticing that 9 of 11 colonies had changed sex, we conducted another two rounds of sampling in March and April 2011 not only to re-confirm the sex change phenomenon, but also to rule out the possibility of sampling error. The analysis showed that as of February 2011, the sex of 9 of 11 colonies was different from that in 2010 **(Fig. 2 A)**. In 2 out of 11 colonies (#6 and 7), the sex remained female **(Fig. 2 A)**.

**Figure 2.**
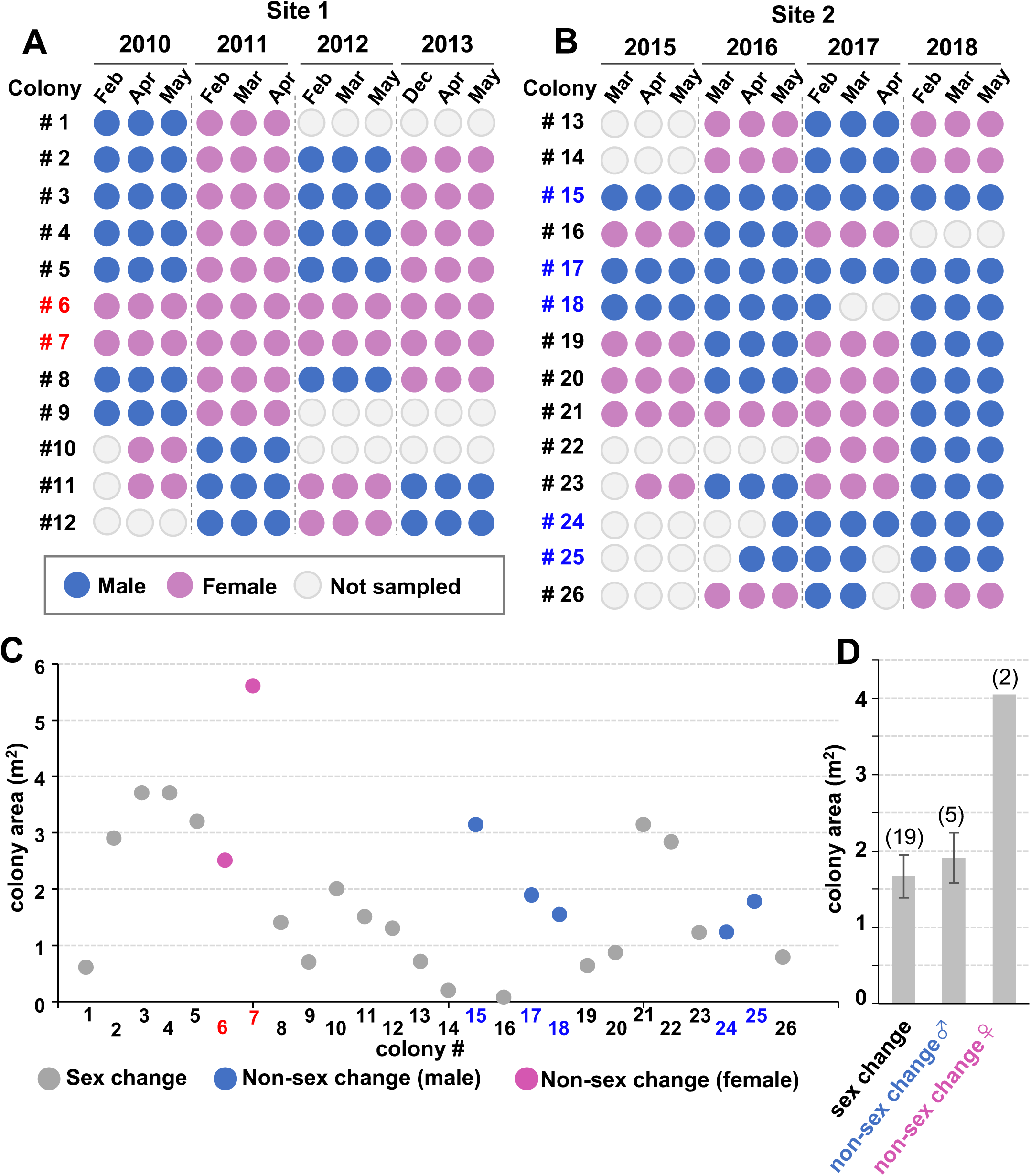
Sexes of *F. ancora* colonies and their size. **A** and **B**. Sexes of colonies at Site 1 (**A**) and Site 2 (**B**). At Sites 1 and 2, surveys were performed during 2010-2013 and 2015-2018, respectively. In each survey year, fragments of each colony were sampled at different months and their sexes were histologically determined. Each circle indicates the result of histological analysis. Results are shown in blue for males and in pink for females. In cases in which sampling was not possible due to colony mortality, inclement weather, etc., circles are indicated in grey. **C**. Sizes (area, m^2^) of colonies analyzed in this study. Sex-change colonies are shown in grey, non-sex-change male colonies are shown in blue, and non-sex-change female colonies are shown in pink. **D.** Comparison of colony width between sex-change colonies and non-sex-change colonies. Numbers in parentheses indicate numbers of colonies.

In February 2012, we resampled the labeled colonies. Three of the labeled colonies (# 1, 9, and 11) were dead and could not be sampled. However, 9 labeled colonies (8 colonies labeled in 2010, 1 colony newly labeled in 2011) were resampled three times in 2012. The analysis showed that the sexes of colonies # 2, 3, 4, 5, 8, and 11 had switched back to the original sexes in 2010 **(Fig. 2 A)**. Two colonies #6 and 7 remained female.

The following year, from December to May 2013, we resampled the labeled colonies 3 times and performed the same analysis, which again confirmed the sex change phenomenon in 7 of 9 colonies, with 2 colonies remaining female **(Fig. 2 A)**.

Subsequently, we conducted similar surveys from 2015-2018 for a total of 14 new colonies at Site 2 which was located about 100 m from Site 1. As at Site 1, there were both annual sex-change colonies and non-sex-change colonies **(Fig. 2 B)**. At Site 2, five non-sex-change male colonies were identified. Non-sex-change female colonies were not found at Site 2.

Through 8 years of observation, we examined 26 colonies at Sites 1 and 2 (**Table 1**). Among them, 19 colonies (73.1%) changed their sex every year, while the remaining 7 colonies (26.9%) were fixed as male or female. The number of non-sex-change female and non-sex-change male colonies were 2 and 5 (16.7% and 35.7% of the total), respectively.

**Table 1.** Summary of *F. ancora* colonies during 2010-2013 and 2015-2018.

| Sampling site | Period | No. of colony | Annual sex | Non-sex change |
| --- | --- | --- | --- | --- |
|  |  | examined | change (%) | (%) |
| Site 1 | 2010-2013 | 12 | 10 (83.3 %) | 2 (16.7 %) females |
| Site 2 | 2015-2018 | 14 | 9 (64.3 %) | 5 (35.7 %) males |
| Site 1 + 2 | (total of 8 years) | 26 | 19 (73.1 %) | 7 (26.9 %) females<br>& males |

### Location and colony size of sex-change and non-sex-change colonies

Next, we investigated whether there were any differences in growth environment and colony size between sex-change and non-sex-change colonies. At both Sites 1 and 2, sex-change and non-sex-change colonies were located within 1-10 m (**Fig. 1 F and G**), and there appeared to be no environmental differences. No significant differences were observed in colony size (**Fig. 2 C and D**). There were also no prominent differences in colony color, as seen underwater. Most were either green or light green.

### Timing and processes of sex change

Subsequently, we sought to discover the process and timing of sex change by histological analysis. Samples collected during two reproductive seasons, from several months before spawning, through a few weeks after spawning, to the next spawning season were examined.

First, the process of female-to-male sex change was investigated (**Table 2**). All gonads collected during the female phase (December to May, **Fig. 3 A**) were ovaries filled with many oocytes. No testicular tissues, such as male germ-cell clusters, were observed in the ovaries (n=136, **Fig. 3 A, Table 2**). Analysis of gonads collected 0-3 months after spawning (May-August, n=52) showed that about 80% were sexually undifferentiated gonads with undifferentiated germ cells (**Fig. 3 A and B**). The remaining ∼20% were gonads with residual or newly developed small oocytes (**Fig. 3 A and C, Table 2**). However, further analysis revealed that oocyte numbers decreased with time after spawning (**Fig. 3 D)**. Two months after spawning (July), about 50 oocytes were found in a gonad, but by 3 months post-spawning (August), the number decreased to about 10 oocytes (**Fig. 3 D)**. Five months post-spawning (October), no oocytes were observed in the gonads, and initiation of spermatogenesis was observed (n=147, **Fig. 3 D, Table 2**).

**Figure 3.**
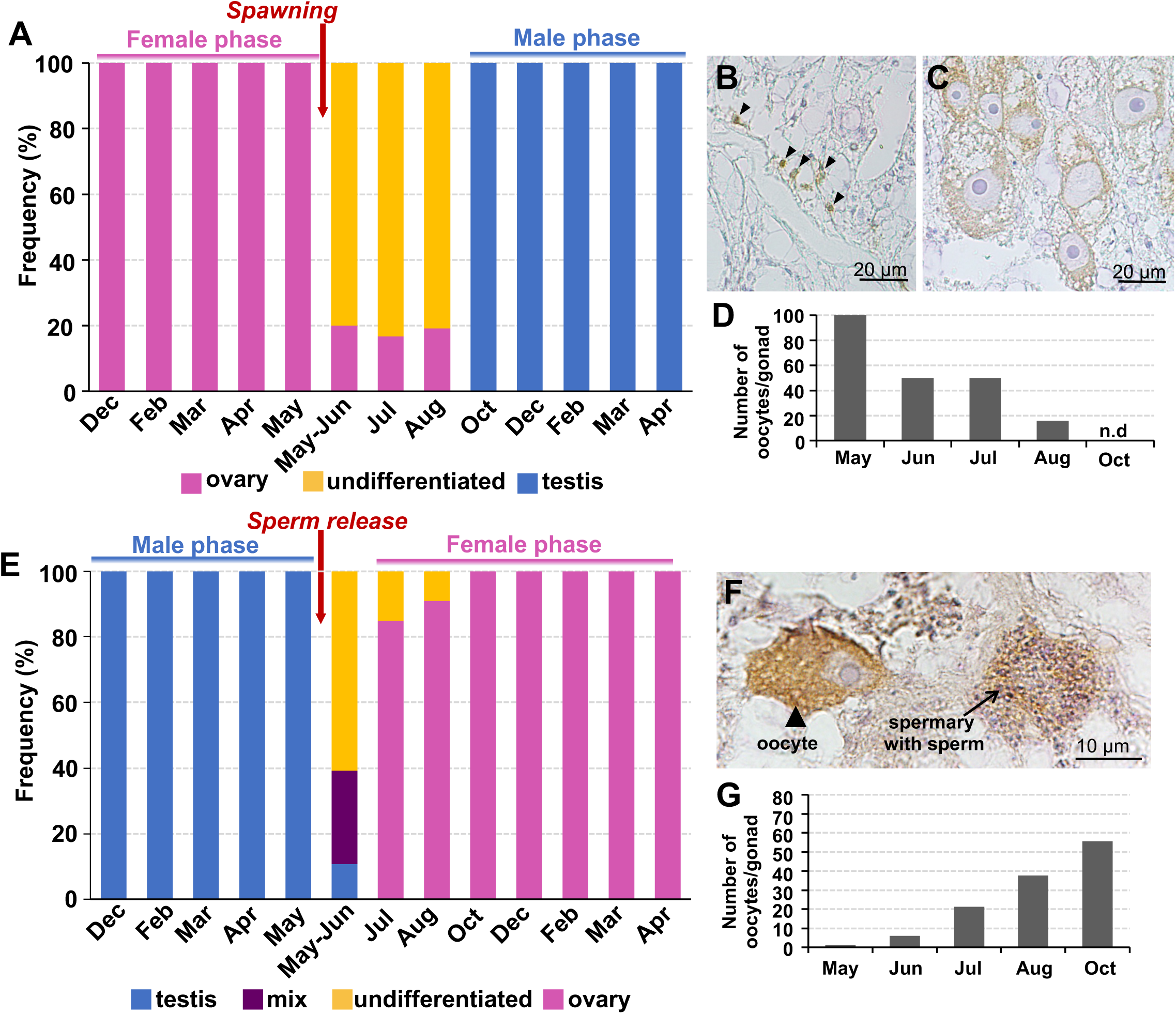
Timing and processes of sex change in *F. ancora*. **A-D**. The process of sex change from female to male. **A**. Observed types of gonads and their frequencies in each month. **B**. Representative immunohistological section of an undifferentiated gonad. Undifferentiated (early-stage) germ cells are labeled brown by immunostaining with an anti-*F. ancora* vasa antibody (arrows). **C**. Representative immunohistological section of an ovary. Oocytes are labeled brown with an anti-*F. ancora* vasa antibody. **D**. Average number of oocytes per gonad. **E-G**. Sex change from male to female. **E**. Observed types of gonads and their frequencies each month. **F**. Representative immunohistological section of a gonad (mix) that has an oocyte and residual sperm (spermary with sperm). Oocytes and sperm are labeled brown with an anti-*F. ancora* vasa antibody and an anti-*F. ancora* rGC antibody, respectively. **G**. Average numbers of oocytes per gonad.

**Table 2.** Results of histological analyses of female-to-male sex change.

|  | No. of<br>colony<br>investigated | No. of gonad<br>investigated | No. of<br>gonad<br>with<br>spermaries | No. of<br>gonad with<br>oocytes | No. of gonad<br>with both<br>spermary/sperm<br>and oocytes | No. of sexually<br>undifferentiated<br>gonad |
| --- | --- | --- | --- | --- | --- | --- |
| **Dec** | **8** | **38** | **0** | **38** | **0** | **0** |
| **Feb** | **6** | **28** | **0** | **28** | **0** | **0** |
| **Mar** | **3** | **15** | **0** | **15** | **0** | **0** |
| **Apr** | **6** | **42** | **0** | **42** | **0** | **0** |
| **May** |  |  |  |  |  |  |
| **(before spawning)** | **4** | **13** | **0** | **13** | **0** | **0** |
| **May-Jun** |  |  |  |  |  |  |
| **(after spawning)** | **4** | **20** | **0** | **4** | **0** | **16** |
| **Jul** | **2** | **6** | **0** | **1** | **0** | **5** |
| **Aug** | **6** | **26** | **0** | **5** | **0** | **21** |
| **Oct** | **3** | **8** | **0** | **0** | **0** | **8** |
| **Dec** | **6** | **21** | **21** | **0** | **0** | **0** |
| **Feb** | **9** | **36** | **36** | **0** | **0** | **0** |
| **Mar** | **7** | **15** | **15** | **0** | **0** | **0** |
| **Apr** | **7** | **67** | **67** | **0** | **0** | **0** |

Next, the process of male-to-female sex change was investigated. All gonads collected during the male phase (December-May, **Fig. 3 E**) were testes filled with spermaries. No oocytes were observed in testes (n=167, **Fig. 3 E, Table 3**). Analysis of gonads collected a few weeks after sperm release (May-June, n=56) revealed that about 60% were sexually undifferentiated gonads with undifferentiated germ cells, and that about 30% were gonads having small numbers of both newly developed small oocytes and residual sperm (**Fig. 3 F, Table 3**). Three months after sperm release (August), more oocytes were observed in about 90% of the gonads (**Fig. 3 E and G**, n=67, **Table 3**). Further analysis revealed that oocyte numbers increased as time passed from the month of sperm release **(Fig. 3 G)**. Active oogenesis with apparent oocyte growth was also observed.

**Table 3.** Results of histological analyses of male-to-female sex change.

|  | No. of colony<br>investigated | No. of gonad<br>investigated | No. of<br>gonad with<br>spermaries | No. of<br>gonad with<br>oocytes | No. of gonad<br>with both<br>spermary/sperm<br>and oocytes | No. of sexually<br>undifferentiated<br>gonad |
| --- | --- | --- | --- | --- | --- | --- |
| **Dec** | **6** | **21** | **21** | **0** | **0** | **0** |
| **Feb** | **9** | **36** | **36** | **0** | **0** | **0** |
| **Mar** | **7** | **15** | **15** | **0** | **0** | **0** |
| **Apr** | **7** | **67** | **67** | **0** | **0** | **0** |
| **May** |  |  |  |  |  |  |
| **(before** | **6** | **28** | **28** | **0** | **0** | **0** |
| **spawning)** |  |  |  |  |  |  |
| **May-Jun** |  |  |  |  |  |  |
| **(after** | **10** | **56** | **6** | **0** | **16** | **34** |
| **spawning)** |  |  |  |  |  |  |
| **Jul** | **4** | **23** | **0** | **20** | **0** | **3** |
| **Aug** | **7** | **44** | **0** | **40** | **0** | **4** |
| **Oct** | **6** | **11** | **0** | **11** | **0** | **0** |
| **Dec** | **8** | **38** | **0** | **38** | **0** | **0** |
| **Feb** | **6** | **28** | **0** | **28** | **0** | **0** |
| **Mar** | **3** | **15** | **0** | **15** | **0** | **0** |
| **Apr** | **6** | **42** | **0** | **42** | **0** | **0** |

## Discussions

The present study revealed a unique reproductive strategy of *F. ancora*. Of 26 colonies we examined, about 70% changed their sex every year, while the remaining 30% did not. Although sex change has been reported in various taxa (1-5, 18-20), to the best of our knowledge, there is no description of organisms such as *F. ancora*, which change sex every year. *Fimbriaphyllia ancora* will be an ideal model for studying mechanisms of sex differentiation and plasticity, and reproductive ecology.

Although sex-change has been reported in some fungiid corals (10-12), the timing and process of sex change, as well as biological characteristics of those corals have not been explored. The present study revealed histologically when and how sex-change occurred in *F. ancora* (**Fig. 4 A and B**). In female-to-male sex change, small oocytes were still observed in some gonads up to about 3 months after spawning (late May to Aug), but none were observed 5 months after spawning (Oct). This suggests that corals were in an intermediate phase or female phase for 0-3 months after spawning, and that female-to-male sex change occurred 4-5 months after spawning. In contrast, in male-to-female sex change, the earliest oocytes appeared several weeks after sperm release (late May-June). Oocytes appeared in most gonads 3 months after sperm release (Aug). This suggests that male-to-female sex change occurred from 0-3 months after sperm release.

**Figure 4.**
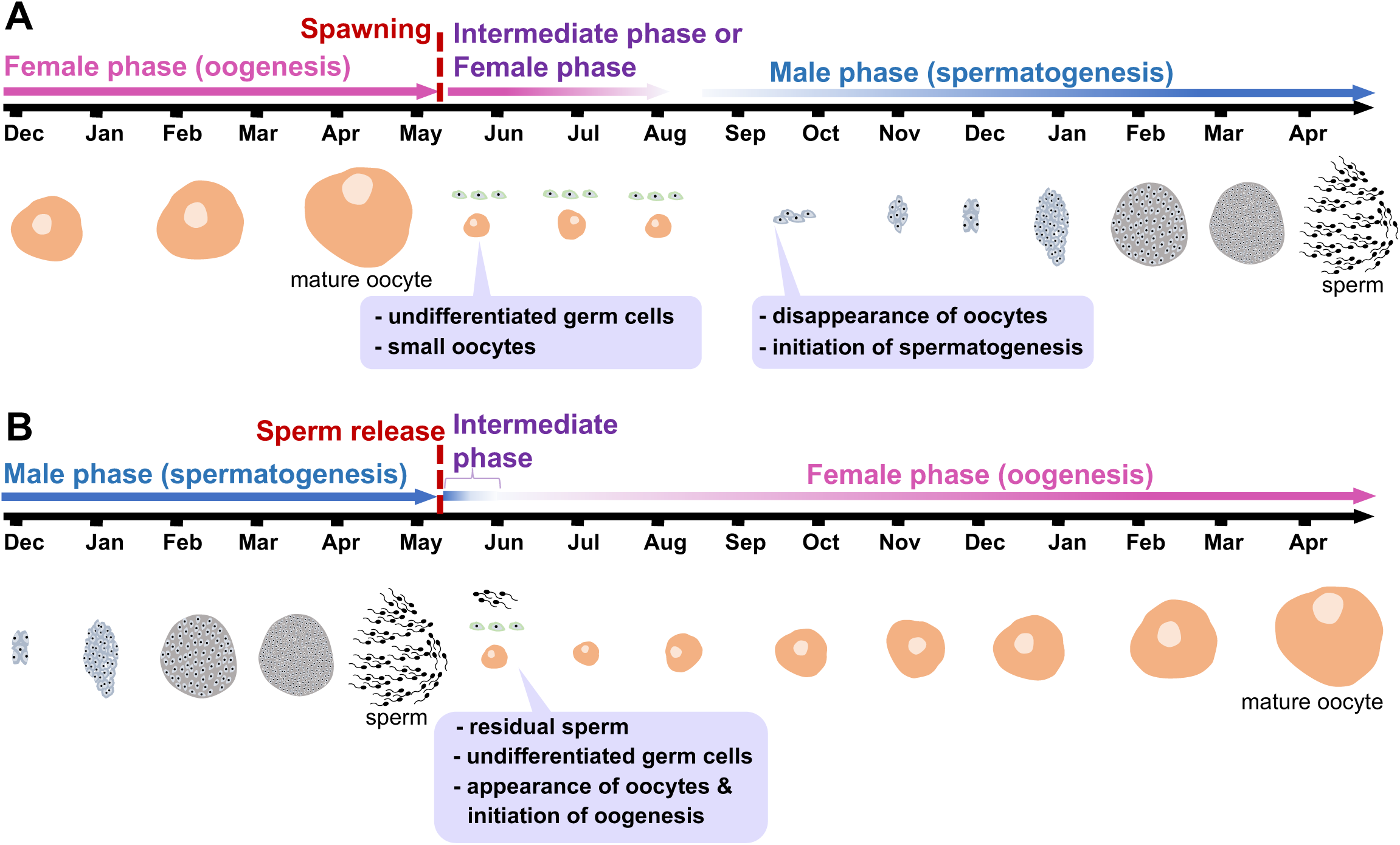
Schematic diagram of possible processes of sex change in *F. ancora*. **A**. Female-to-male sex change. From 0-3 months after spawning (late May-August), colonies still do not change completely into males, and small oocytes and undifferentiated germ cells are present in the gonads (intermediate phase or female phase). Five months after spawning (October), oocytes are no longer present and the male phase (spermatogenesis) was initiated. **B**. Male-to-female sex change. The earliest oocytes appeared a few weeks after sperm release (late May-June), and residual sperm, undifferentiated germ cells, and small oocytes were observed in some gonads (intermediate phase). Oocytes appeared in most gonads 2-3 months after sperm release (July-August) and the female phase (oogenesis) was initiated.

Sex change phenomena have been well studied in fish (3,5,21). Based on differences in biological characteristics, organisms that change sex can be further classified into two types: (i) organisms that have both testes and ovaries simultaneously. Testes or ovaries are selectively developed according to their sexual phase. For example, in the bi-directional sex-changing gobiid fish, *Trimma okinawae*, individuals in the female phase have well-developed ovaries, but inactive testes. Conversely, individuals in the male phase have mature testes, but immature ovaries (5, 22); and (ii) organisms that produce eggs or sperm in the same gonad according to their sexual phase. For example, during female-to-male sex change in the grouper, *Epinephelus coioides*, ovarian tissue degenerates and disappears completely, while testicular tissue appears and develops, i.e., gonads completely transform from ovary to testis (23, 24). In *F. ancora*, in the process of male-to-female sex change, we found that residual sperm and oocytes coexisted transiently in the gonads shortly after sperm release. Then, the gonads eventually developed into ovaries in a few months. This finding suggests that *F. ancora* is more like the second type.

In *Hydra oligactis*, which belongs to the same phylum as corals, females can emerge from male clones, particularly when animals are cultured at high temperatures (25). In *F. ancora*, however, sex-change and non-sex-change colonies are found as little as 1 m apart. Thus, it is unlikely that the growing environment of these colonies is different, suggesting that sex change in *F. ancora* is unrelated to environmental factors such as temperature, light, flow, etc.

Previous reports of sex change in corals have linked individual/colony size and sex change. For example, in the stony coral, *Stylophora pistillata*, small colonies are male, but once they exceed a threshold size, they become simultaneous hermaphrodites (26). Protandrous sex change has been reported in some non-sessile corals belonging to the Family Fungiidae, and there is a relationship between individual/colony size and sex change (10,11). In the case of *F. ancora*, on the other hand, no differences in colony size were observed between sex-change and non-sex-change colonies. There were also no noticeable differences in appearance, e.g., colour and morphology of tentacles, among them. These results suggest that in *F. ancora*, sex change may be related to intrinsic factors, e.g., age and genetics. Future detailed investigations of ages and genomes in relation to sex-change and non-sex-change colonies will clarify this hypothesis.

In *Hydra*, it has been demonstrated that the sex of an individual polyp is determined by sex-specific germline stem cells (GSCs) in the polyp (27). If a polyp has male GSCs, which can only produce male germ cells, the polyp becomes male. The same applies to females. If we apply this *Hydra* sex-determination system to *F. ancora*, the following hypothesis can be considered. Non-sex-change male or female *F. ancora* colonies may contain only male-GSCs or female-GSCs. On the other hand, in the case of sex-change colonies, GSCs of both sexes may be present in every gonad. In the male phase, gonads are functionally testes. However, a small number of female GSCs may also be present, and they undergo self-renewal to increase in number. Soon after, when most sperm are released during spawning, female GSCs, which are predominant in the gonad after sperm release, differentiate into oocytes and eventually, the gonad becomes an ovary. Proliferation of male GSCs in the testis during spermatogenesis may be repressed by some undetermined mechanism. The same applies to the change from female to male. In *Hydra*, molecular markers for female GSCs and male GSCs have been identified (28, 29). In our previous study, we have identified GCS-like cells in gonads of *F. ancora* (30). Identification of molecular markers for female GSCs and male GSCs in *F. ancora* will allow us to test the hypothesis.

*Fimbriaphyllia ancora* was functionally gonochoric at each reproductive season. However, long-term studies across multiple reproductive seasons in this study demonstrated that many colonies can produce gametes of both sexes, i.e., that they are sequential hermaphrodites. In 2012, some colonies of the stony coral, *Diploastrea heliopore,* were reported to change sex after the reproductive season (31). However, since only 5 colonies were labeled and studied for 14 months spanning two reproductive seasons, it is currently unknown whether there are colonies that change their sex annually. Nevertheless, our findings and this previous report suggest that corals described as gonochoric may include species undergoing sex change. Determining the reproductive mode of corals will require labeling colonies and studying them through multiple reproductive cycles.

What advantages are there in having both annual sex-change colonies and non-sex-change colonies within a population? One possible advantage would be that the presence of the both increases the probability of successful sexual reproduction. Because most corals are sessile, they are unable to pursue mating partners. This disadvantage may be mitigated if there are sex-change colonies in the population. The presence of an annual sex-change colony near a non-sex-change male or female colony would give them a chance to reproduce sexually at least once every two years with a high probability.

## Summary

We discovered that many colonies in the stony coral, *F. ancora* change sex annually. We also showed that female-to-male sex change occurs 4-5 months after spawning, whereas male-to-female sex change occurs 0-3 months after sperm release.

## Materials and Methods

### Collection of F. ancora

To collect *F. ancora*, we selected 2 sites in Nanwan Bay, southern Taiwan (Site 1, 21°57’205N, 120°46’116E; Site 2, 21°57’221N, 120°46’092E) because of accessibility and ease of anchoring. Species identification of *F. ancora* was performed based on two external characteristics, flabello-meandroid skeletons and anchor-like tentacles (32). *Fimbriaphyllia ancora* colonies were labeled with numbers, and distances between colonies were determined by scuba divers using a measuring tape. For sampling, portions of labeled colonies (approximately 5-8 cm/fragments) were collected at different times by scuba divers using a hammer and chisel during the reproductive season in 2010-2013 for site 1 and 2015-2018 for site 2. To avoid collecting young polyps that lacked gonads, central parts of colonies were selected and sampled. Furthermore, when resampling was performed within a month or two, a region at least 10 cm away from the previously sampled location was selected. To reveal the process and timing of sex change, sampling was performed at Sites 1 and 2 during 16 months spanning two reproductive seasons, 1∼5 months before spawning (in December, February, March, and April), a few weeks before and after spawning (in May and June), and 1-11 months after spawning (in July, August, October, December, February, March, and April). Collecting was approved by the administration of Kenting National Park (Permit Number: 1010006545). All collected *F. ancora* fragments were fixed for 16 h at 4°C in Zinc Formal-Fixx (Thermo Shandon, Pittsburgh, PA), diluted 1:5 with 0.22 μm-filtered artificial seawater. After decalcification with 5% formic acid, samples were preserved in 70% ethanol until use.

## Histological analysis

Fixed samples were embedded in paraplast (Thermo Scientific), and serial 4-µm sections were prepared with a microtome (Thermo Shandon, Pittsburgh, USA). Since approximately 100-600 serial sections constituted an entire gonad, all gonadal sections were collected. Then, sections were de-waxed, hydrated, and stained with Hematoxylin and Eosin Y (H & E staining, Thermo Shandon). The number of oocytes in a gonad was determined by observing all sections of each ovary under a microscope (BX51; Olympus). All analyses were performed with Image J software (Wayne Rasband, National Institutes of Health, Bethesda, MD, USA; https://imagej.nih.gov/ij).

## Immunohistochemistry

Sample fixation, decalcification, embedding, and sectioning were performed according to methodologies described above. Hydrated sections were incubated for 30 min with HistoVT ONE (Nacalai Tesque, Inc, Kyoto, Japan) for antigen retrieval, and then incubated for 10 min in 3% H2O2, and blocking for 1 h in 5% skim milk. Sections were then incubated for 16 h at 4 °C in anti-*F. ancora* vasa antibody (vasa: germ cell marker (33) diluted 1:4,000 in phosphate-buffed saline containing 0.1% Tween 20 [PBT] with 1% skim milk. For detecting sperm, sections were incubated with anti-*F. ancora* receptor guanylate cyclase (rGC) antibody (rGC: sperm marker) (34) diluted 1:1,000 in PBT with 1% skim milk for 16 h at 4 °C. After washing three times with PBT, sections were incubated with a biotinylated goat anti-rabbit IgG antibody (Vector Laboratories, Burlingame, USA; diluted 1: 4,000 in PBT with 2% skim milk) for 30 min. Sections were then incubated in avidin-biotin-peroxidase complex (ABC) solution (Vector Laboratories), and the immunoreactivity (brown colour) was visualized using 3, 3’-diaminobenzidine (DAB, Sigma-Aldrich) system. Immunostained sections were then counter-stained with Haematoxylin (blue-purple colour), and photographed under a microscope (BX51; Olympus).

## Statistics

All data are presented as means ± standard deviations (SDs) or standard error (SE). Statistical significance was determined using one-way ANOVA followed by Tukey’s test with a statistical significance level of P<0.05. All analyses were performed using Statistical Package for the Social Sciences (SPSS) software.

## Ethics

Experiments were carried out in accordance with principles and procedures approved by the Institutional Animal Care and Use Committee, National Taiwan Ocean University.

## Author contributions

SS conceived and designed the experiments. YLC, PHT, and SS performed the experiments. YLC, PHT, and SS analyzed the data. SS wrote the manuscript. CFC modified the manuscript.

## Acknowledgments

We are grateful to all the lab members who assisted us in collecting samples and measuring the colonies and distances between colonies. We also thank Dr. Steven Aird for his great help in preparing the manuscript.

